# Regional history outweighs phylogeny in shaping recent demographic trajectories of Indo-West Pacific sea snakes

**DOI:** 10.64898/2026.10.11.757437

**Authors:** Amelia Rose Pointon, Alastair John Ludington, Vinay Udyawer, Vhon Oliver Garcia, James Nankivell, Balazs Buzas, Balazs Farkas, Csaba Geczy, Kate Laura Sanders

## Abstract

**Aim:** Understanding how biodiversity is formed and distributed is fundamental to anticipating how populations will respond to ongoing climate change and conserving future diversity. Sea snakes are a recent marine radiation comprising replicate lineages separated by vicariance and long-term isolation among major Indo-Pacific ocean basins. This system provides a powerful opportunity for testing whether demographic trajectories are shaped by regional history, phylogeny, or geographic range size. We therefore compare the demographic and phylogeographic histories of fully marine sea snakes to evaluate the role of Pleistocene climate change in structuring biodiversity across major biogeographic regions, including the global marine hotspot of the Indo-Australian Archipelago.

**Location:** Indo-Pacific tropical and sub-tropical marine.

**Methods:** We quantified demographic trajectories using whole-genome sequencing and integrated these with a broad-scale phylogeographic analysis of genome-wide single nucleotide polymorphisms (SNP). SNP sequencing used 652 individuals to investigate the phylogeographic structure of 20 species across the Indian Ocean, Southeast Asian and Australian West-Pacific biogeographic regions. Whole genome data from 36 individuals provided coalescent estimates of effective population size (*N_e_*) through time. These *N_e_* trajectories were then analysed using phylogenetic comparative methods to test the effects of regional history, phylogeny, and geographic range size.

**Results:** Distinct within-species phylogeographic structure was uncovered between the regions supporting vicariance across biogeographic barriers. *N_e_* trajectories in the last 200 thousand years (kyr) were more strongly influenced by regional history than by phylogeny or contemporary range size. Southeast Asian lineages maintained the largest historical populations and experienced the shallowest population declines compared to lineages in the Indian Ocean and Australian West-Pacific. These results are consistent with the role of the marine biodiversity hotspot centred on the Indo-Australian Archipelago (IAA) as a region of long-term population persistence and stability during Pleistocene climatic fluctuations.

**Main Conclusions:** Demographic responses to Pleistocene climate change are strongly region-specific across the Indo-West Pacific, reflecting variation in habitat persistence and connectivity among ocean basins. Over the last 200kyr, regional biogeographic history is a stronger determinant of demographic history than shared ancestry and species range size. Based on these patterns, we recommend a hierarchical conservation framework in which ocean basins represent primary management units, and within-basin genetic structure informs finer-scale spatial planning. Of particular concern are Indian Ocean lineages, which exhibit lower effective population sizes and face intense fishing pressure and habitat loss. In contrast, Australian populations maintain relatively high genetic diversity and may represent an important regional refuge under future change. The demographic stability of Southeast Asian lineages supports the IAA as a long-term centre of population persistence, but this historical resilience offers little protection against current pressures: shallow marine habitats in the region are heavily degraded and sea snakes are directly harvested for trade.

## 1 Introduction

Over the past 2.6 million years, repeated glacial cycles have reshaped the geographic distribution of marine biodiversity, through fragmentation and expansion of populations over broad spatial scales. Understanding how populations have responded to these climatic fluctuations, and whether these responses have been geographically consistent across species’ ranges, is fundamental to interpreting present day patterns of biodiversity and predicting vulnerability to ongoing climate change. However, for most taxa and regions, this long-term demographic context remains poorly resolved, particularly outside well studied continental terrestrial systems.

This is especially important for widely distributed marine groups whose populations span oceanographic regions with distinct climatic and geological histories. Exceptional marine biodiversity is centered on the Indo-Australian Archipelago (IAA), spanning the tropical waters of Southeast Asia from the Andaman Sea to the western Pacific (Barber and Bellwood, 2005; Bellwood et al., 2005; Cowman and Bellwood, 2013; Hughes et al., 2002; Yasuhara et al., 2022). At its core, the Coral Triangle holds the highest marine biodiversity on Earth and is characterised by structurally complex seascapes and a mosaic of deep-water barriers that have alternately promoted isolation and connectivity among populations across glacial cycles (Bellwood et al., 2012). During Pleistocene glacial maxima, sea levels reductions of up to ∼120 m exposed continental shelves and contracted shallow marine habitat, while simultaneously creating a network of relatively stable habitat refugia that promoted population persistence (Voris, 2000). Interglacial transgression re-submerged the Sunda and Sahul shelves, creating a vast expanse of shallow habitat and reconnecting previously isolated marine populations. This alternation of stability and fragmentation is thought to have promoted both long-term persistence and the accumulation of lineage diversity in the IAA relative to surrounding marine biogeographic regions in the Australia-Pacific and Indian Ocean.

Adjacent ocean regions differ markedly from the Coral Triangle and broader Indo-Australian Archipelago (IAA). The Indian Ocean is geologically young, and its eastern margin against the Sunda Shelf is a tectonically active, providing limited opportunity for shelf formation. Much of Indian subcontinent has large sediment outflows due in part to the erosion of the Himalayas, but this is a geologically recent scenario. The Indian Ocean basin therefore has narrower continental shelves and less bathymetric complexity than the IAA, and correspondingly fewer opportunities for stable refugia during glacial cycles (Harris et al., 2014; Thomas et al., 2024). Northern Australia and subtropical and tropical adjacent areas support expansive shallow continental shelves but much of the northern and eastern shelf was exposed during the Last Glacial Maximum, concentrating shallow tropical seas on the north-west shelf, while the island-arc terranes of Melanesia provide little shelf development (Heap and Harris, 2008; Wilson et al., 2013). These regional contrasts in climatic and environmental history predict divergent demographic trajectories: IAA lineages should maintain larger and more stable effective population sizes through time, while peripheral populations should show greater instability and more frequent bottlenecks.

Sea snakes (Hydrophiinae) provide an unusually tractable system for quantifying regional differences in demographic trajectories across co-distributed taxa. These fully marine reptiles are a recent, species-rich radiation, with 62 recognised species diversifying across the Indo-West Pacific within the last few million years (Sanders et al., 2013a; Sanders et al., 2013b). Importantly, multiple closely related sea snake lineages are co-distributed in shallow-water continental shelf habitats in the Australian, Indian Ocean, and Southeast Asian biogeographic regions, and mitochondrial and microsatellite markers show that most widespread species comprise reciprocally monophyletic intraspecific lineages corresponding to ocean basin, with narrowly distributed species showing basin-level endemism (Buzas et al., 2019; Ukuwela et al., 2014, 2016, 2017). These shared signals of Pleistocene vicariance, replicated across regions within a single radiation, make sea snakes a tractable system for asking whether demographic trajectories reflect regional environmental history or species-specific dynamics.

Coalescent analyses of whole diploid genomes provide an opportunity to reconstruct changes in effective population size (N_e_) through time. N_e_ is a key parameter in conservation genetics; it reflects the number of individuals effectively contributing genes to future generations and provides an indication of long-term genetic diversity and demographic history (Nadachowska-Brzyska et al., 2022). High N_e_ values generally indicate large or historically stable populations and can also result from the reconnection or admixture of previously divergent subpopulations, whereas low N_e_ values may indicate population decline, bottlenecks or long-term small population size and can be associated with reduced genetic diversity and adaptive capacity.

The conservation status of most sea snakes remains poorly understood due to a lack of knowledge on species distributions, ecology, population trends, and taxonomic status (Udyawer et al., 2018). The most recently published IUCN Red List assessment in 2010, classified 34% of sea snakes as data-deficient, preventing assessment of their threat status (Elfes et al., 2013). The principal threats are habitat loss, commercial fishing, coastal development and climate change. Fishing pressure has been linked to population declines and shifts in regional assemblages in India and the Gulf of Thailand (Cao et al., 2014; Rao, 2021), and short-range endemics in both Southeast Asia and Australia are vulnerable to localised disturbance (Elfes et al., 2013; Rasmussen et al., 2012). As ectotherms with metabolically demanding diving habits, sea snakes are also expected to be sensitive to rising ocean temperatures (Udyawer et al., 2016).

In this paper, we quantify historical demographic trajectories of co-distributed sea snakes using whole genome sequencing (WGS) for 36 individuals across 18 species. We complement this analysis with the first broad-scale phylogeographic analysis of sea snakes across their Indo-West Pacific range, using high-resolution genome-wide single nucleotide polymorphism (SNP) data for 652 individuals across 18 localities and 20 species. Our findings provide new insights into how marine biodiversity has been shaped by regional differences in climatic and evolutionary history during the Pleistocene.

## 2 Methods

### 2.1 Samples

Tissue samples were sourced from 656 individuals from 20 species across the global distribution of sea snakes in the Indian Ocean (IO), Southeast Asia (SEA) and the Australia-West-Pacific (AUS) (Figure 1). These were collected between 2002 and 2024 by the authors and their collaborators, or sourced from museum collections (Museum and Art Gallery of the Northern Territory, MAGNT; South Australian Museum, SAM; Western Australian Museum, WAM; Queensland Museum, QM; Museum Zoologicum Bogoriense, MZB; Field Museum of Natural History, FMNH). Liver, skin, vertebra and blood samples were obtained under appropriate animal ethics approval and scientific collection permits held by the collector. Of the 652 samples used for SNP genotyping, 425 were sequenced for this study, 156 were from Coppersmith et al. (in prep) and 71 from Nankivell & Sanders (2025); a further five samples were used for whole-genome resequencing (Supplementary Table S1 and S2).

**Figure 1.**
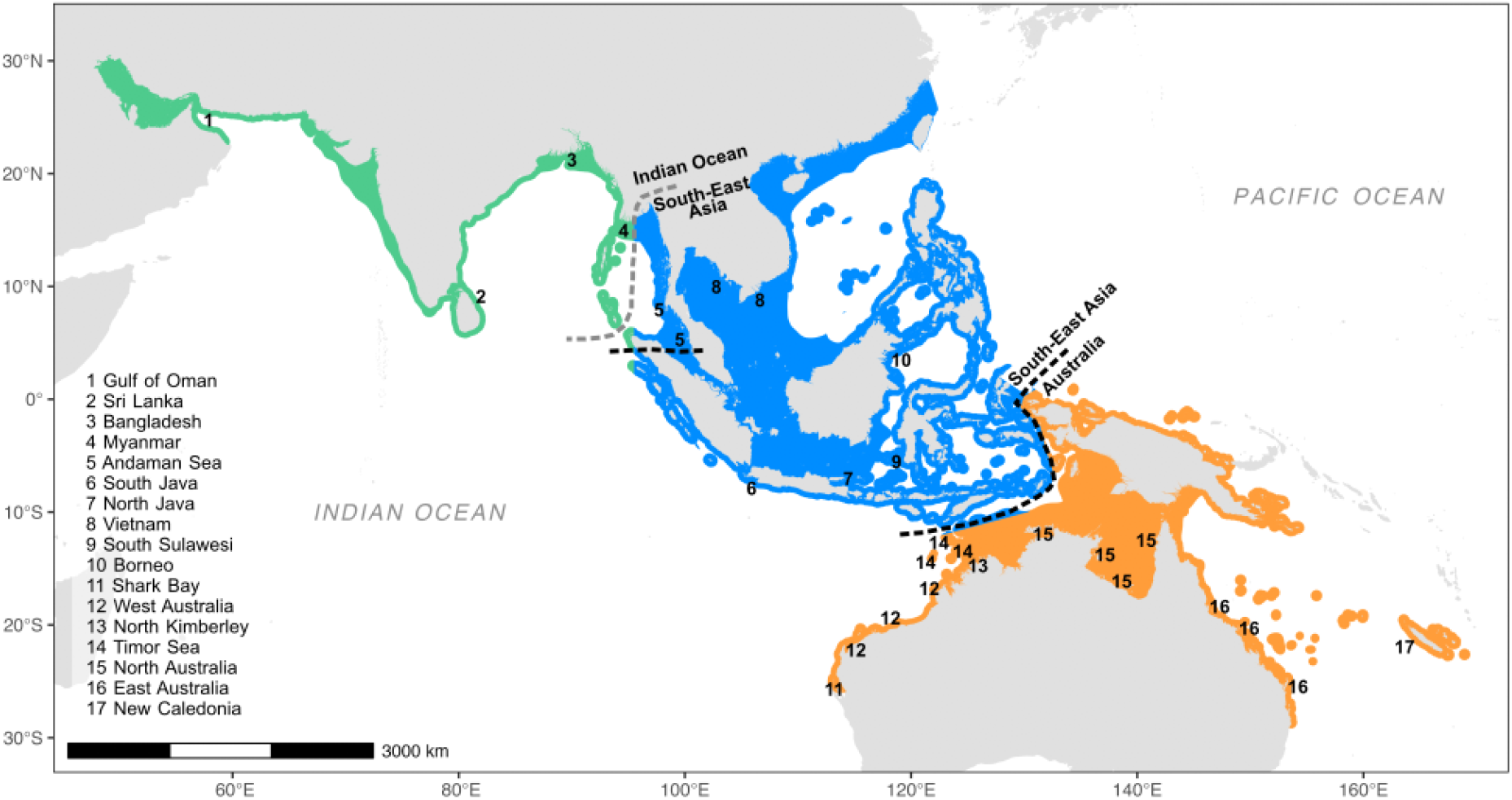
Global distribution of sea snakes within the three biogeographic regions; Indian Ocean (green), Southeast Asia (blue) and Australia (orange). Numbers indicate localities sampled in this study. Dashed lines indicate phylogeographic breaks between basins. The grey dashed line marks the alternative break between the Indian Ocean and South Asia for *Hydrophis curtus* and *Hydrophis schistosus*.

### 2.2 DNA extraction and sequencing

Genomic DNA was extracted using the Gentra Puregene tissue kit (Gentra 2014) following the manufacturer’s protocol with three modifications: (1) tissue was ground directly in cell lysis solution; (2) an in-house lysis buffer (10 mM Tris/2% SDS/0.1 M EDTA) replaced the kit lysis solution; and (3) ammonium acetate replaced the protein precipitation solution. DNA was quantified using a Quantus fluorometer, with samples required to yield >500 ng at 50 ng/μl. Samples were concentrated using a Centrivap DNA concentrator where necessary and quality-checked by gel electrophoresis. DNA was then sent to Diversity Arrays Technology Pty Ltd (DArTseq, Canberra, Australia) for SNP genotyping using a proprietary genome-complexity reduction pipeline with PstI and HpaII restriction enzymes [28,29]. Following digestion, ligation and amplification, samples were sequenced across eight sequencing runs on an Illumina HiSeq 2500 between 2020 and 2024.

Short-read whole genome sequences were produced for five individuals at the Ramaciotti Centre for Genomics using the Illumina DNA PCR-Free library preparation kit and sequenced on an S4 flow cell with paired-end 150bp sequencing with an Illumina NovaSeq 6000. Combined with 29 whole genomes previously generated by the authors (Ludington et al., 2023; Rossetto et al., 2024) and two genomes obtained from GenBank (Li et al., 2021), the final set comprised 22 lineages from Australia, nine from Southeast Asia and five from the Indian Ocean (detailed in Supplementary Table S2).

### 2.3 Sequence data QC and variant calling

All SNP datasets were passed through the same quality control pipeline: adapters and cut-site overhang removed from demultiplexed sequencing reads using the bbmap function in bbduk (Bushnell 2024); contaminant microbial and human DNA sequences removed with Kraken2 v2.1.2 using the standard database (Wood et al., 2019); and low quality bases (--average_qual 10) and reads below a minimum length (--length_required 25) removed using fastp v.0.23.2 (Chen, 2023). Each quality control step was visualised in multiqc (Ewels, 2016).

Three types of SNP datasets were generated: genus-level sets (*Hydrophis– Microcephalophis*; *Aipysurus–Emydocephalus*) for phylogenetic inference; a set of representative lineages matching the whole-genome samples for phylogenetic signal testing; and per-species sets containing all available populations for population genetic analysis. Where the same individual was not available for DArT sequencing, an individual from the same population was substituted. Per-species SNP sets were generated using all available populations for population genetic analysis. All SNP calling used the de novo approach in ipyrad v0.9.93 (Eaton and Overcast 2020) on the Adelaide University High-Performance Computing (HPC) system, Phoenix. Default parameters were set except for stricter adapter filtering (=2) and to trim reads to 69 bases long where newer sequencing runs contained a longer read resulting in batch effects. All post-run analysis was performed on the outputted Variant Call Format (VCFs) files. The sequencing replicate samples with the lowest sample coverage were removed from the VCFs. Phylogenetic SNP sets were further filtered in BCFtools (Danecek et al., 2021) with outliers removed (-S), SNPs filtered for Minor Allele Frequency (MAF>=0.01) and missingness (F_MISSING<=0.25), and one random SNP per loci kept (+prune -w 100 -n1). Species SNP sets for population genetics were processed in snpR (Hemstrom and Jones, 2023) using R Statistical Software (v4.5.1, R Core Team 2021). Unless otherwise stated, each VCF was filtered to keep only SNPs with a MAF greater than 0.01 (maf = 0.01) and present in 90% of samples (min_ind = 0.9) and with one random SNP kept per RAD loci. See Supplementary Table S3 for sequence and SNP-calling statistics.

Short-read whole genome sequences were mapped to the respective phylogenetically closest lineage of seven available high-quality reference genomes. Samples were aligned to autosomal sequences of the respective reference genomes using BWA-mem2 (Vasimuddin et al., 2019), except for the *Aipysurus laevis* genome (GCA_040207615.1) which was first modified to remove Z chromosome sequences. Z Chromosome contigs were identified by mapping the *A. laevis* genome to the *H*. *major* reference genome (GCA_033807585.1) using minimap2 (li 2018). Alignment statistics were calculated using *rustybam stats* (Vollger and Hulselmans, 2023) and contigs removed if more than 50% of their length was aligned to the Z chromosome. Our samples had an average alignment coverage of 30% ranging from 13% to 85% (Supplementary Table S4). Mappability files for each reference genome were then generated using GenMap v1.3.0 (Pockrandt et al., 2020). Input files for demographic inference in MSMC2 were then produced from the alignment files using the accessory scripts *bamCaller.py* and *generate_multihetsep.py*. Variant calling was performed separately for each chromosome and sample using BCFtools mpileup v1.15.1 (-q 30 -Q 30 -C 50) followed by BCFtools *call* (-c). The resulting variant calls were processed with *bamCaller.py* to generate per-chromosome VCF and mask files. Together with the mappability files, these were then passed to *generate_multihetsep.py* to produce formatted inputs for MSMC2.

### 2.4 Population structure -Tree building

Before tree building, VCFs of phylogenetic SNP sets were converted to phylip files using vcf2phylip.py v2.0 (Ortiz, 2023). To investigate the phylogenetic history of the species of interest we built separate maximum likelihood phylogenetic trees of the *Hydrophis-Microcephalophis* and *Aipysurus-Emydocephalus* clades with IQTree3 (Thomas et al., 2026). Ascertainment bias correction for SNP data (GTR+ASC) was performed with the discrete gamma model (+G) and ultrafast bootstrap approximation (-B 1000, --nmax 2000). As expected, IQTREE returned an error for invariant sites and produced an output file for each dataset containing only variant sites (*.varsites*). These were then used as the input data for final tree building using 532 individuals and 648 SNPs for the *Hydrophis-Microcephalophis* tree and 120 individuals and 5,610 SNPs for the *Aipysurus-Emydocephalus* tree. Both trees were rooted post hoc on the node connecting sister genera *Aipysurus-Emydocephalus* and *Hydrophis-Microcephalophis*.

A separate maximum likelihood phylogeny was inferred using SNPs for the whole-genome representatives in RAxML version 7.2.8 using the GTRGAMMA substitution model (Stamatakis 2014). To account for ascertainment bias in the in SNP-only data, the ASC_GTRGAMMA model was applied. The rapid bootstrap algorithm (-f a) was used to simultaneously perform tree inference and bootstrap resampling. Statistical support was assessed using 100 rapid bootstrap replicates. The resulting tree was used for downstream comparative analyses.

### 2.5 Population structure – PCA, admixture, and diversity metrics

Population structure was assessed for each per-species SNP set using Principal Component Analysis (PCA) in the R package snpR (Hemstrom & Jones 2023) and an ‘admixture’ like estimation of ancestry proportion with fastMIXTURE (Santander et al., 2024), excluding species with less than three localities or samples (*Aipysurus fuscus* and *Microcephalophis gracilis*). Individuals were grouped by sampling locality to visualise genetic clusters between populations and ocean basins. To perform fastMIXTURE, clean VCFs were first converted to plink BED format with pLink2 (Chen et al., 2019). Ancestry proportion was estimated for a range of clusters (*K*) informed from the PCAs and tested with 5-fold cross-validation (--cv 5) (Supplementary Figure 1). Optimal *K* values were chosen where the graphed error rate dropped and additional ancestral groups provided only marginal improvement. Two further species, *A. foliosquama* and *Hydrophis kingii* were excluded due to Eigenvalues not converging due to too few SNPs. Adjacent sampling localities with no genetic structure were then grouped to represent populations for calculating genetic diversity metrics.

To compare patterns of genetic diversity across ocean basins we used the snpR function, get.snpR.stats to calculate the weighted mean Nucleotide Diversity (Hemstrom and Jones, 2023), Observed Heterozygosity (H_o_) and Tajima’s *D* per population. For H_o_ estimation, we used a missing data threshold of 0% to reduce biases in small sample sizes following recommendations by Schmidt et al. (2021). In Tajima’s D estimation, the Minor Allele threshold was not set to meet the assumptions of the analysis. We only considered populations with greater than or equal to five individuals for comparisons across ocean basins.

### 2.6 MSMC2 analysis of demographic history

We reconstructed the historical trajectories of effective population size (*N*_e_) across the three ocean basins using short-read whole-genome sequencing using MSMC2 (Schiffels & Wang, 2020). MSMC2 was run twice, first using the default clock (1*2+25*1+1*2+1*3) then with a custom clock (2*2+25*1+2*3). Both runs were performed with 100 EM iterations (-I 100) and 20 bootstrap replicates (See Supplementary Figure 2 for raw *N*_e_ curves with bootstraps). MSMC2 outputs were then processed and visualised in R. Exploratory visualisation of the demographic curves revealed no significant difference between the clocks therefore the default was used in all further analysis. We set a y-axis maximum limit of 1e6 after exploratory visualisation revealed an optimum resolution of demographic signal. MSMC2 curves were scaled to years using a mutation rate calculated using the known divergence time of the outgroup genus, *Microcephalophis*, at 12 million years ago (Sanders et al., 2016). We applied Jukes-Cantor correction to the proportion of SNPs from total sites called, scaled by the divergence time and generation time of five years (Kishida et al., 2025) for a mutation rate of 6.45e-09 per site/per generation (following Ludington and Sanders, in prep). After an initial assessment of the raw MSMC curves, a minimum time boundary of 20kya and maximum time boundary of 3MYA was applied where MSMC modelling produced informative curves. An outlier sample, SRR11461820, was removed from further analysis. In all further analysis we considered a maximum time boundary of 1.5MYA to represent time intervals after the inferred divergence of sea snakes, and a minimum time boundary of 50kya to remove inflated *N*_e_ estimates in recent time intervals. To compare patterns in overall demographic fluctuations across ocean basins, we followed the methods of Germain et al. (2023) and first interpolated the N_e_ trajectory of each sample at 121 comparative equidistant log-linear time intervals. Each lineage was assigned to an ocean basin validated by the genetic structuring of each species in PCA and fastMIXTURE analysis. We then calculated the average N_e_ at each interval within species per basin before averaging *N_e_* per basin and performing min-max normalisation to rescale the values between 0 and 1.

To enable investigation of phylogenetic signal in *N*_e_ trajectories, we applied the previous interpolation of sample-specific *N*_e_ trajectories onto a common set of time points. We chose 146 timepoints spaced every 10 thousand years between 50kya and 1.5kya. *N_e_*estimates at each time point along this standardised vector were then log10-transformed and ordered from the most recent to the oldest prior to analysis of phylogenetic signal and phylogenetic analyses of relationships between *N*_e_, ocean basin and geographic range size. Each lineage was therefore represented as a multivariate trait vector of 146 log10-transformed *N*_e_ values. Subsets of this vector corresponding to specific time windows were used in subsequent analyses. We assigned each lineage to one of the three defined biogeographic regions: Australian (AUS), the Indian Ocean (IO), and Southeast Asia (SEA); based on phylogenetic and population genetic structure analysis.

The maximum-likelihood phylogeny estimated for all 36 lineages using RAxML was used to construct a phylogenetic variance–covariance (VCV) matrix, which defined the covariance structure among lineages for all subsequent comparative analyses under a Brownian motion model of trait evolution.

### 2.7 Phylogenetic signal in N_e_

Whether effective population size (*N*_e_) histories are more similar among closely related taxa than expected by chance was tested by quantifying phylogenetic signal in the log_10_ *N*_e_ vector using the multivariate K statistic ‘Kmult’ (Adams, 2014). Kmult is an extension of Blomberg’s K that measures phylogenetic signal in high-dimensional traits by comparing observed among-species similarity to that expected under a Brownian motion model of evolution along the phylogeny. To examine how phylogenetic signal changed through time, we calculated Kmult across sequential time windows of 50,000 years spanning from 50,000 to 500,000 years ago. Statistical significance was assessed using 999 tip-label permutations. All phylogenetic signal analyses were conducted in R v4.6.0 using the geomorph package (Baken et al., 2021) and Kmult was extracted by the physignal() function for each time window (Adams et al., 2026).

### 2.8 Phylogenetic MANOVA of N_e_ among ocean basins

A phylogenetic multivariate analysis of variance (phylogenetic MANOVA) was performed to test whether log-transformed *N*_e_ trajectories of sea snake lineages differed across ocean basins, after accounting for shared evolutionary history. A phylogenetic generalised least squares regression framework was used with randomisation of residuals. Statistical significance was assessed using 999 RRPP permutations, with p-values calculated as the proportion of permuted F values greater than or equal to the observed F. Effect size was reported as R², which represents the proportion of variation in *N*_e_ explained by region. Analyses were repeated across time windows of 50,000 years spanning from 50,000 to 500,000 years ago for N_e_ trajectories among the three regions (AUS: n = 22, IO: n = 5 and SEA: n = 9). Because Southeast Asia accounted for most between-group variance in the group comparisons, an additional analysis compared Southeast Asia (n = 9) to outlying regions of Indian Ocean plus Australia (n=27).

Multiple testing across time intervals was corrected for using a permutation-based maxT procedure. For each permutation, the test statistics for all intervals were calculated and the maximum value was used to build a null distribution. Adjusted p-values were then derived by comparing observed statistics to this distribution, accounting for dependence among overlapping intervals. All analyses were conducted in R v4.6.0 using the RRPP package with phylogenetic relationships included via the phylogenetic covariance matrix using lm.rrpp (Collyer and Adams, 2014; Collyer and Adams, 2024).

### 2.9 PGLS regression of N_e_ and range size

To test whether range size (extent of occurrence; EOO) predicts effective population size (*N*_e_) trajectories after accounting for ocean basin, we fitted phylogenetic generalised least squares (PGLS) regression models within a RRPP framework using the phylogenetic variance– covariance structure derived from the RAxML maximum-likelihood tree. The response variable was the log10-transformed *N*_e_ trajectory summarised across 50kya to 150kya time window. This window was selected because it shows strong variance in *N*_e_ among ocean basins (see results).

To produce present day range size estimates for each lineage, we used preliminary extent of occurrence (EOO) datasets developed during the 2026 IUCN Red List of Threatened Species assessment of sea snakes (*in prep*). For lineages occurring in multiple basins, a subset of their distribution map was obtained with each subset based on the biogeographic region groupings defined in population genetic structure analyses. Area was calculated using st_area function of the sf package (Pebesma, 2018) in R. Two models were fitted, the first tested the relationship between *N*_e_ and range size alone (*N*_e_ ∼ log(EOO)). The second model tested the effect of EOO after accounting for ocean basin (*N*_e_ ∼ region + log(EOO)), with Type I sequential sums of squares such that the effect of EOO represents variation in *N*_e_ not explained by region. The Southeast Asian *Hydrophis schistosus* lineage was excluded due to uncertainty in range boundaries for this clade, resulting in a sample size of n = 35. Both *N*_e_ and EOO were log-transformed prior to analysis.

To construct the phylogenetic variance–covariance matrix, we used the R package, ape (Paradis and Schliep, 2019). PGLS models were fitted using the RRPP package with lm.rrpp(). Statistical significance was assessed using permutation-based *F*-statistics and associated *p*-values from the RRPP framework.

## 3. Results

### 3.1 Phylogeographic patterns: trees, clustering, and ancestry proportions

Phylogeographic patterns were assessed for each species in the MSMC2 analyses using a combination of maximum-likelihood tree building, Principal Component Analysis (PCA) and ancestry proportion inference (fastMIXTURE). PCA and fastMIXTURE rely on allele frequency covariance and are more robust to missing data, whereas concatenated SNP datasets represent an ascertainment-biased subset of variable sites and frequently contain uneven missingness that violates the assumptions of substitution models used in likelihood phylogenetic inference. For these reasons the concatenated SNP trees are likely to resolve shallow population structure less reliably, and we therefore emphasise the patterns recovered by PCA (Figure 2) and fastMIXTURE (Figure 3) over those in the SNP trees (Figure 4).

**Figure 2.**
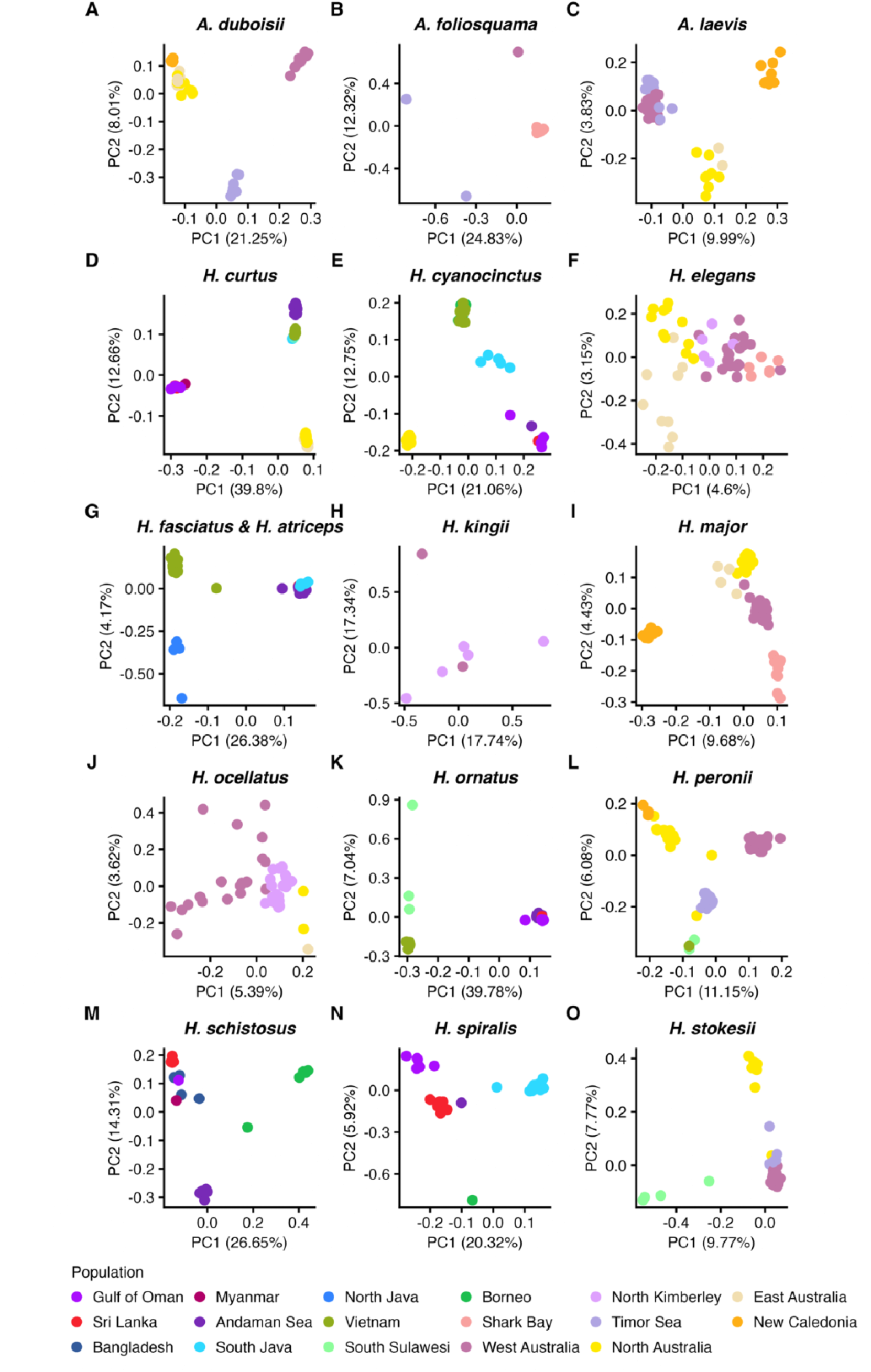
Principal Component Analyses of 16 sea snake species sampled across their Indo-Pacific distributions. Each individual is represented by a point coloured by population. The percentage of total variance explained by the Principal Component is displayed in axes.

**Figure 3.**
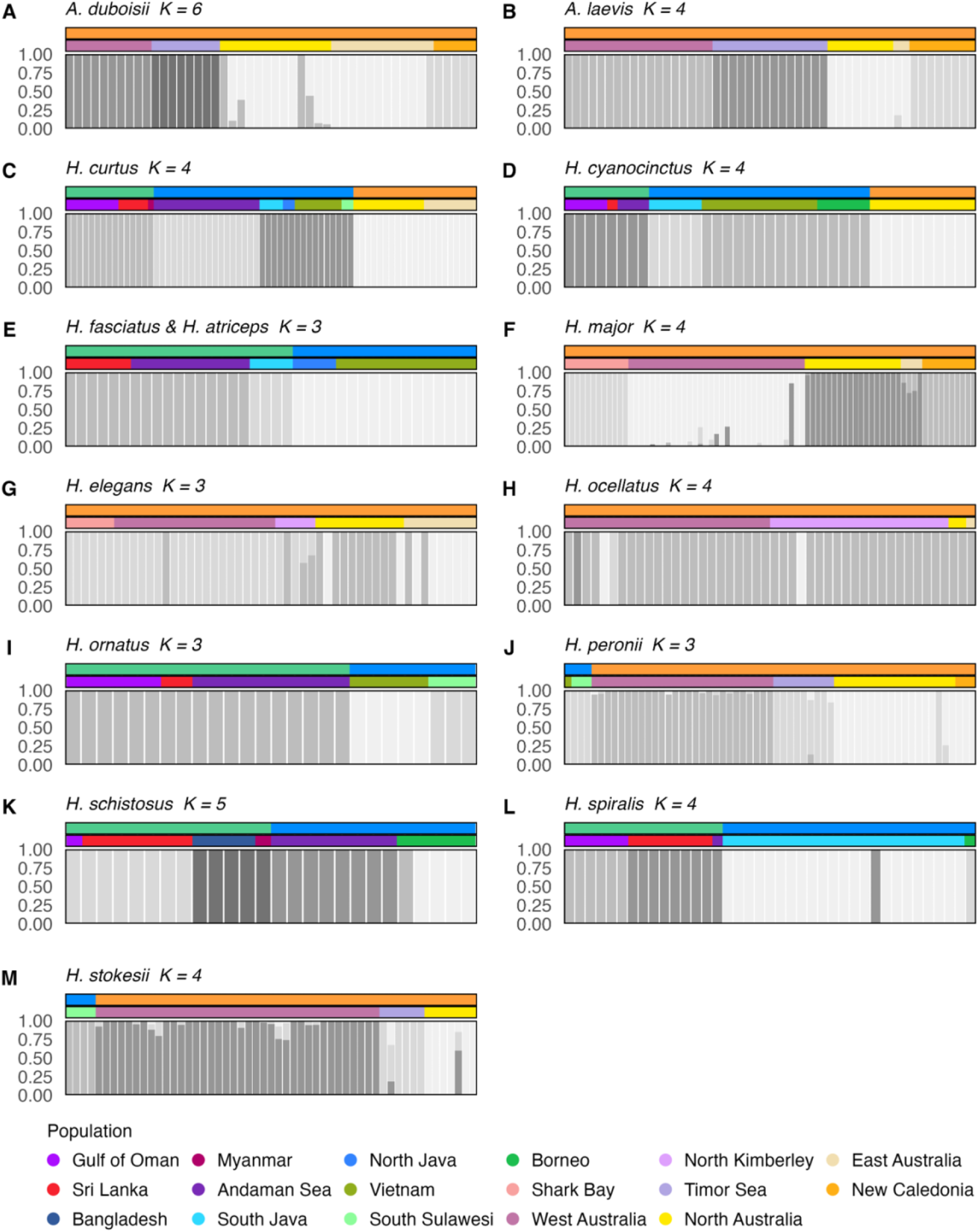
Ancestry proportion estimates (fastMIXTURE) for 14 species of sea snakes. Individuals (grey vertical bars) are coloured by the proportion of ancestry assigned to an ancestral cluster (*K*). Individuals are ordered by population geography from west to east. Metadata is displayed above each plot with population in the middle panel and region in the top panel (IO = green, SEA = blue, AUS = orange).

**Figure 4.**
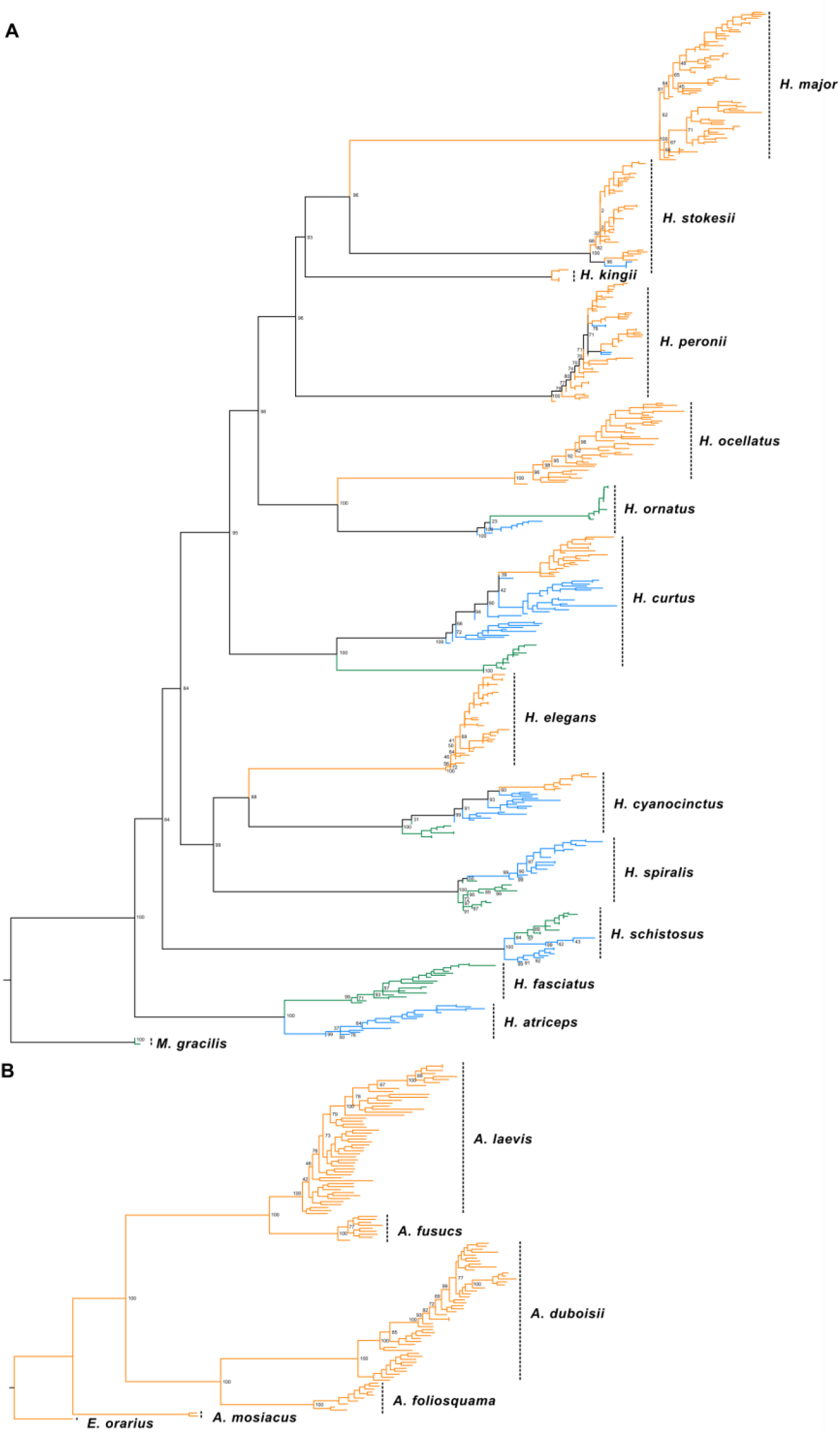
Maximum Likelihood SNP tree (IQTREE) of the Indo-West Pacific distribution of sea snake species with corresponding whole-genome sequence. Separate trees represent (A) *Hydrophis-Microcephalophis* species (n = 15) and (B) *Aipysurus-Emydocephalus* species (n = 6). Trees were midpoint-rooted using nodes representing the common ancestors of *Hydrophis* and *Microcephalophis,* and *Aipysurus* and *Emydocephalus*. Branch lengths are proportional to nucleotide substitutions per site. Clades are coloured by ocean basin grouping (AUS = orange, SEA = blue, IO = green) following parsimonious inference.

Of the 14 *Hydrophis* species represented in the tree, seven had distributions and sampling spanning multiple ocean basins. Between four to 70 individuals across one to ten populations were sampled per species (Table 1). For structure analysis, the number of SNPs after filtering for each species ranged from 5246 to 1252 SNPs.

**Table 1.** Per species SNP dataset for Indo-West Pacific sea snakes used in population genetic structure and diversity statistical analyses.

| Genus | Species | Individuals | Populations | SNPs (Structure and Nucleotide Diversity) | SNPs (Observed Heterozygosity) | SNPs (Tajima's D) |
| --- | --- | --- | --- | --- | --- | --- |
| *Aipysurus* | *duboisii* | 47 | 5 | 2491 | 545 | 2491 |
| *Aipysurus* | *foliosquama* | 10 | 3 | 2492 | 902 | 2492 |
| *Aipysurus* | *fuscus* | 10 | 1 | 5246 | 3601 | 5246 |
| *Aipysurus* | *laevis* | 50 | 5 | 4240 | 514 | 4240 |
| *Hydrophis* | *curtus* | 70 | 10 | 1763 | 21 | 2809 |
| *Hydrophis* | *cyanocinctus* | 39 | 7 | 2363 | 359 | 2363 |
| *Hydrophis* | *elegans* | 50 | 5 | 2532 | 177 | 2532 |
| *Hydrophis* | *fasciatus & atriceps* | 38 | 5 | 3272 | 399 | 3272 |
| *Hydrophis* | *kingii* | 7 | 2 | 4272 | 4272 | 4272 |
| *Hydrophis* | *major* | 76 | 5 | 2521 | 199 | 4102 |
| *Hydrophis* | *ocellatus* | 42 | 4 | 2143 | 39 | 2143 |
| *Hydrophis* | *ornatus* | 26 | 5 | 1252 | 229 | 1252 |
| *Hydrophis* | *peronii* | 61 | 6 | 1699 | 33 | 3184 |
| *Hydrophis* | *schistosus* | 26 | 6 | 1640 | 180 | 1640 |
| *Hydrophis* | *spiralis* | 38 | 5 | 2604 | 240 | 2604 |
| *Hydrophis* | *stokesii* | 55 | 4 | 2864 | 309 | 4377 |
| *Microcephalophis* | *gracilis* | 4 | 2 | 1921 | 1921 | 1921 |

*Hydrophis curtus* was sampled across all three basins, and all analyses recovered strong geographic structuring among them. PC1 (37.94%) separated Indian Ocean localities from Southeast Asia and Australia, and PC2 (10.48%) separated Southeast Asia from Australia. Ancestry assignment recovered four distinct clusters, with 0% shared ancestry between basins. In contrast, substantial shared ancestry was found among clusters within basins: 100% between North Australia and East Australia; 100% shared across Indian Ocean populations. Within Southeast Asia, there were two clusters with 0% shared ancestry, one for Andaman Sea and another between North Java, South Java, Vietnam and South Sulawesi populations. The tree recovered the same deepest split, between an Indian Ocean clade and Southeast Asia plus Australia, with maximal bootstrap support, although Australia and Southeast Asia were not reciprocally monophyletic, in contrast to previous mitochondrial and microsatellite studies; Australian samples nonetheless formed a distinct cluster separated by a long branch. Andaman Sea lineages formed their own ancestry cluster but fell within the Southeast Asia clade in both tree and PCA, supporting their assignment to that basin. Together these analyses indicate strong structuring across the Andaman Sea, with Australian lineages isolated more recently. Overall, multivariate PCA, admixture and phylogenetic analyses support strong structuring between Indian Ocean and Southeast Asia biogeographic regions east and west of the Andaman Sea, with Australian lineages isolated more recently, albeit on evolutionary timescales that are meaningful to the present study.

*Hydrophis cyanocinctus* also showed the same three-basin divergence across all analyses (PC1 = 20.33%; no shared ancestry between basins), with South Java being intermediate between Indian Ocean and Southeast Asia; these samples had high missingness and were excluded from the phylogeny. The deepest divergence separated Indian Ocean lineages from the common ancestor of Southeast Asia and Australia, with Australian lineages, previously recognised as *H. pacificus*, forming a distinct cluster nested within the Southeast Asia grouping. All populations east of the Andaman Sea were most closely affiliated with Southeast Asia.

The allopatric sister species *H. fasciatus* and *H. atriceps* were recovered as reciprocally monophyletic groups in the Indian Ocean (*H. fasciatus*) and Southeast Asia (*H. atriceps*), With Southeast Asian populations diverging east of the Andaman Sea. Both ancestry assignment and PCA returned distinct clusters for Indian Ocean and Southeast Asia populations with PC1 accounting for 26.91% of the variance between these basin-endemic species clusters. PC2 (3.92%) revealed within basin structure between Vietnam and North Java in *H. atriceps*, and between South Java and a shared cluster of Sri Lanka and Andaman Sea in *H. fasciatus.* These patterns were also supported by ancestry clustering with 100% assignment to unique ancestry clusters in *H. fasciatus and H. atriceps.* In contrast to other species, the Indian Ocean grouping extends east of the Andaman Sea along the southern edge of the Sunda Shelf to include South Java.

Similarly, closely related species *H. ocellatus* and *H. ornatus* diverged across ocean basin regions, between Australia (*H. ocellatus*) and Indian Ocean/Southeast Asia (*H. ornatus*). In *H. ornatus*, strong divergence between Southeast Asia and the Indian Ocean on PC1 (37.93%) obscured within-region structure in the Indian Ocean. Sri Lanka and the Andaman Sea shared an ancestry cluster (100%), with all other populations distinct, and populations east of the Andaman Sea again affiliated with Southeast Asia. *H. ocellatus* showed weak structure within Australia: PCA clusters overlapped along an east–west gradient (PC1 = 5.18%, PC2 = 3.58%) and ancestry was shared across all populations with no geographic structuring.

In *H. schistosus*, which is absent in Australia, Indian Ocean lineages were nested within Southeast Asia lineages, and an early-diverging Andaman Sea clade indicated strong within-basin structure relative to Borneo. PCA separated Indian Ocean samples from two Southeast Asian clusters on PC1 (26.3%) and PC2 (14.59%), with no shared ancestry across basins. Within the Indian Ocean, Gulf of Oman and Sri Lanka formed one ancestry cluster and Bangladesh and Myanmar another.

In *H. spiralis*, the PCA contained strongest clustering between populations with an isolation-by-distance-like pattern emerging across PC1 (21.46%) with populations organised geographically west to east. This trend of population-level structuring was also observed in ancestry assignments with only Sri Lanka and Andaman Sea sharing ancestry clusters (100% assignment). In the tree, a clade of Southeast Asia emerged from the Indian Ocean lineages.

*H. stokesii* was sampled extensively in Australia but only from one Southeast Asia locality (South Sulawesi). Despite this imbalance, Southeast Asian samples formed a clade nested within the Australian lineages. PCA separated Australia from Southeast Asia on PC1 (9.86%) and North Australia from West Australia and the Timor Sea on PC2 (7.72%). All South Sulawesi individuals were assigned entirely to one ancestry cluster; within Australia, 34.21% of Timor Sea and West Australia individuals were admixed (1.68–33.29%), and one North Australian individual carried 82.87% West Australian ancestry.

In *H. peronii,* PCA resolved structure among Australia populations., with Southeast Asia populations central along PC1 (11.06%) and separated from Australia populations along PC2 (6.12%). Both Southeast Asian localities were assigned to a single ancestry cluster that was also represented in the Timor Sea (5.48%), Western Australia (2.29–5.12%) and northern Australia (0.14–2.95%). Within Australia, 47.61% of northern Australian individuals shared 6.29–100% ancestry with New Caledonia, and Western Australia shared ancestry with all other populations (0.60–8.59%). Two Southeast Asian clades were nested within the Australian samples in the phylogeny.

*H. kingii*, *H. major* and *H. elegans* are Australian endemics. *H. kingii* was too sparsely sampled to infer structure, though PCA showed a single overlapping cluster for Western Australia and North Kimberley. *H. major* and *H. elegans* both showed shallow phylogenetic splits and little variance on PC1 (9.39% and 4.44%), indicating weak within-basin structure. In H*. major,* the largest separation was between New Caledonia, which formed a unique ancestry cluster, and all other populations; 35.56% of Shark Bay and other Western Australia individuals shared ancestry (0.58–30.77%), as did 9% of north and east Australian individuals (11.20–100%). In *H. elegans*, Shark Bay and other Western Australian samples formed one cluster, North Kimberley was highly admixed between Western and northern Australia (80% of individuals, 57.57–100%), and eastern Australia formed its own cluster with 15% of individuals fully admixed with northern Australia (the phylogeny did not resolve this moderate structure).

In *A. duboisii*, PCA clustered New Caledonia, East Australia and North Australia, while Timor Sea and Western Australia formed separate clusters along PC1 (18.9%). Ancestry analysis recovered similar structure, with East Australia and North Australia sharing two admixed clusters and the Timor Sea, West Australia and New Caledonia each assigned unique clusters. In the phylogeny, West Australian lineages formed the deepest split, with shallow branches elsewhere and New Caledonian samples forming a divergent cluster nested within North and East Australia.

*Aipysurus laevis* showed structure within Australia but with different groupings: PCA resolved West Australia with the Timor Sea, North with East Australia, and New Caledonia separately, on PC1 (10.21%). Ancestry analysis recovered the same pattern except that Timor Sea individuals formed a distinct cluster from West Australia. The phylogeny again showed shallow branches apart from a nested New Caledonian clade.

The Timor Sea endemic, *A. fuscus*, formed a single phylogenetic clade with shallow branches indicating no genetic sub-structuring. To avoid confounding population diversity estimates, we excluded potential *A. fuscus* × *A. laevis* hybrids that were identified as PCA outliers intermediate to *A. laevis* and *A. fuscus* when graphed together using the *Aipysurus-Emydocephalus* SNP set (and hence removed from the final dataset).

*Aipysurus foliosquama* lineages from Shark Bay and the remainder of West Australia formed a reciprocally monophyletic clade, sister to the Timor Sea lineages. The PCA supports this pattern, with PC1 explaining 25.09% of the variance separating Timor Sea individuals from all conspecifics. Population structure inferences were not made for *A. mosaicus, E. orarius* and *M. gracilis* due to insufficient sampling. Here, structure analysis was used only to recognise outliers and therefore select good candidates for representing the lineage for testing phylogenetic signal in N_e_ trajectories.

Across analyses, the deepest within-species divergences generally coincided with ocean basin boundaries. Reciprocal monophyly between basins was found in *H. fasciatus–H. atriceps* and *H. ocellatus–H. ornatus*, and nested derived clades in *H. cyanocinctus, H. spiralis, H. curtus, H. schistosus, H. stokesii* and *H. peronii*. The position of the Indian Ocean–Southeast Asia break varied among species, falling both east and west of the Andaman Sea. Basal most Southeast Asian lineages in *H. schistosus* and *H. ornatus* indicate an origin in that basin, whereas early-diverging Indian Ocean lineages in *H. curtus, H. cyanocinctus* and *H. spiralis* indicate an Indian Ocean origin, with their deeply nested Australian lineages implying more recent colonisation. Conversely, Southeast Asian lineages of *H. stokesii* and *H. peronii* are nested within Australian clades, supporting an Australian origin for these species.

### 3.2 MSMC2 analysis

Normalised mean effective population size of MSMC2 analyses revealed distinct demographic histories in the three ocean basins (Figure 5). Standard errors of the means overlapped minimally between ocean basins except Indian Ocean and Southeast Asia lineages from 550kya to 240kya. *N*_e_ was highest at the onset of species diversification around ∼1.5-1Mya and was preceded by population declines. At approximately 300kya, *N*_e_ trajectories peaked again in Southeast Asia and Indian Ocean. A comparable second peak in Australia, occurred later at approximately 150kya. Southeast Asia maintains the highest mean *N*_e_ compared to the other basins since the 1.5Mya sea snake diversification. Indian Ocean maintains the lowest *N*_e_ trajectory of all regions for the most recent 200K years. Both Australia and Indian Ocean show increasing *N*_e_ trajectories towards the 50kya margin whereas *N*_e_ in Southeast Asia peaks then decreases again at 60kya. Overlapping standard errors in the most recent time intervals (preceding 50 kya) preclude interpretation of these intervals, although all three basin means trend upwards.

**Figure 5.**
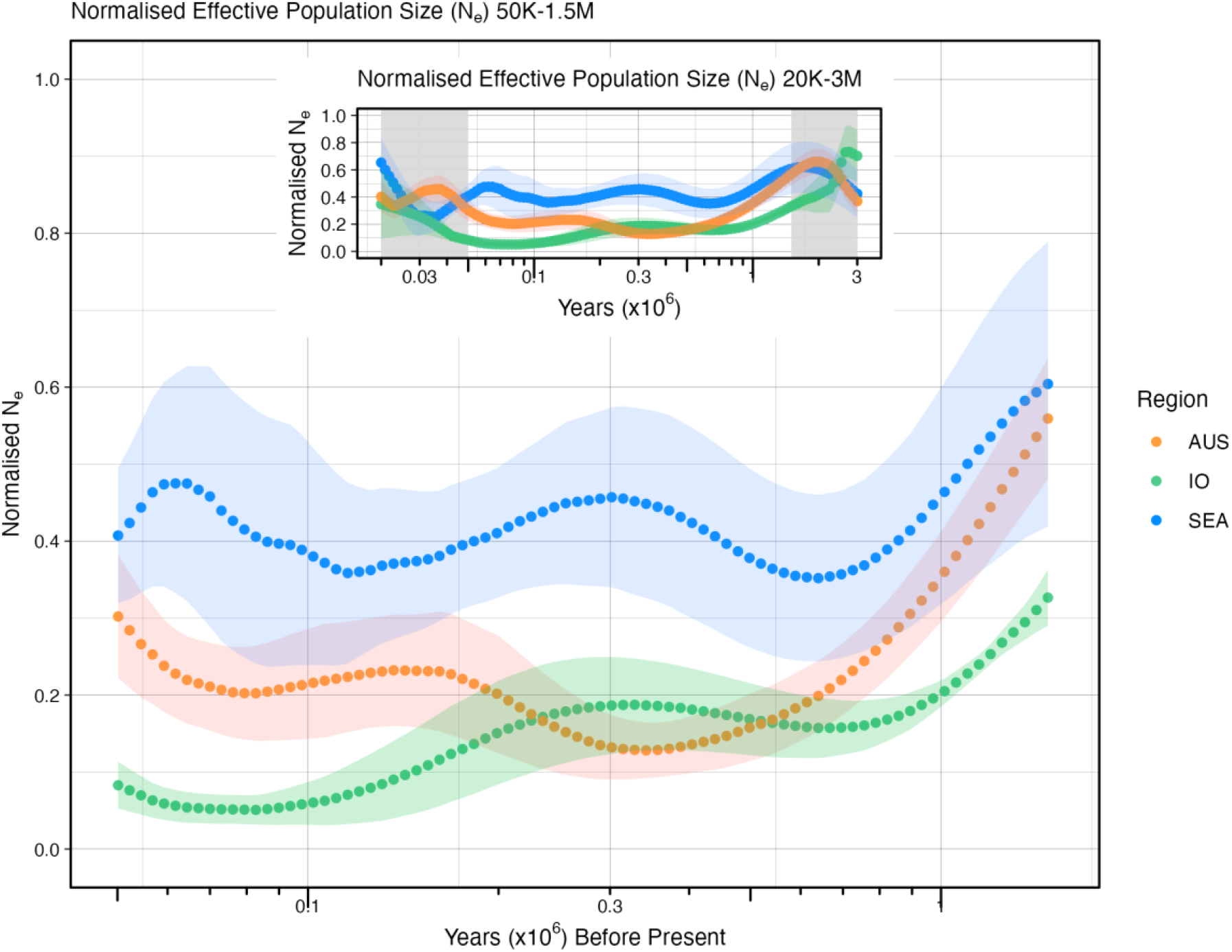
Demographic histories of 36 sea snakes from 50 thousand years ago (kya) to 1.5 million years ago (mya). The extended period from 20 kya to 3 Mya is shown in the inset. The x-axis is presented in the log10 scale. Coloured dots represent the normalised mean N_e_ interpolated for 121 equally spaced (log linear) time points. Ribbon colouring represents the standard error of each mean. Species within Australia are represented in orange, Indian Ocean in green and Southeast Asia (SEA) in blue. The inset shows the full timescale of readable msmc2 curves with grey sections indicating the period of interest.

### 3.3 Phylogenetic signal in *N*_e_ trajectory

To investigate the influence of phylogenetic signal on demographic history, we produced a RAxML maximum likelihood tree comprised of the 36 lineages analysed in MSMC2 using 9,711 SNPs (Figure 6).

**Figure 6.**
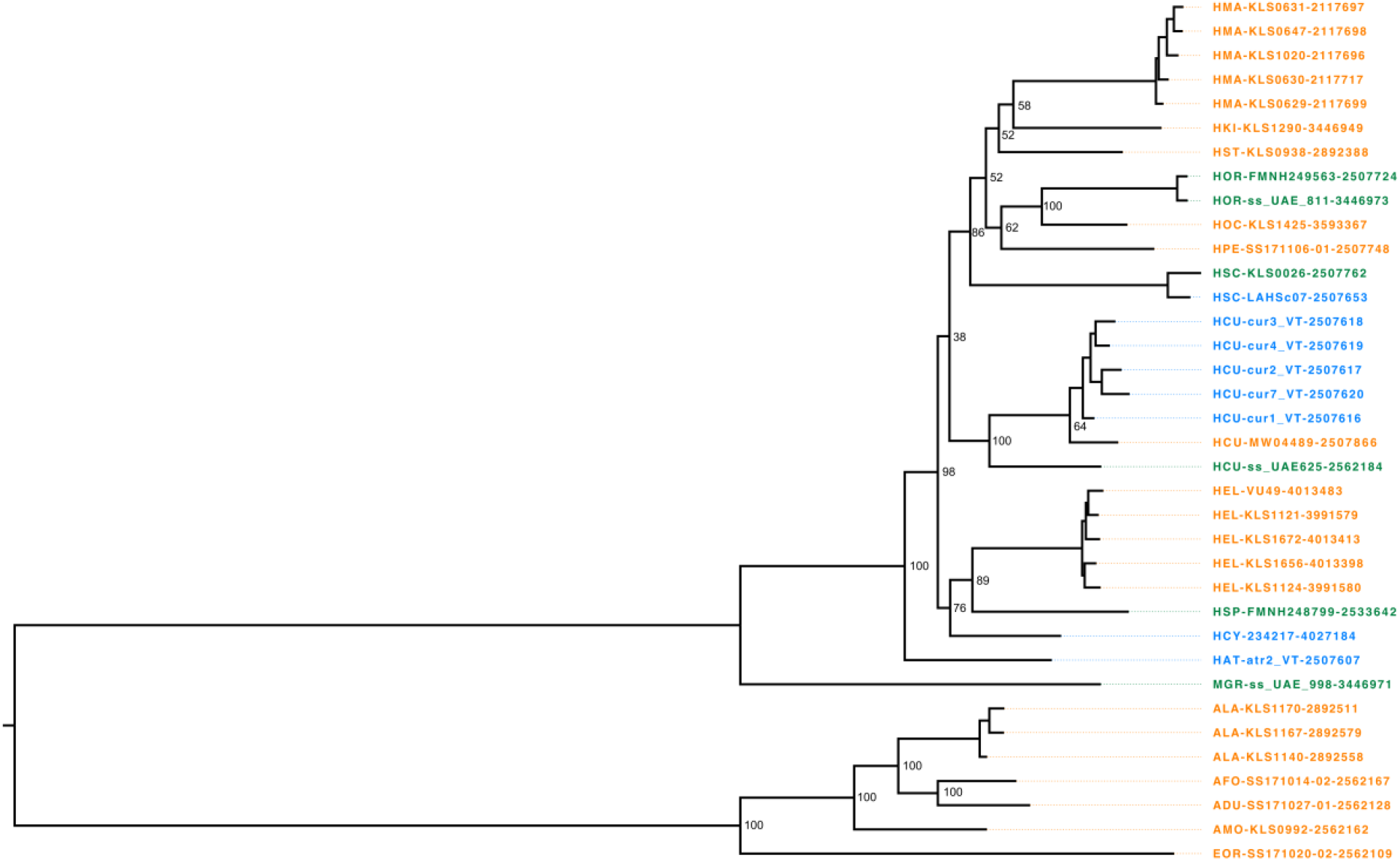
Maximum-likelihood SNP tree showing the phylogenetic relationships of sea snake lineages examined in MSMC2. Lineage names are coloured by ocean basin (Australia-West Pacific = orange, Southeast Asia = blue and Indian Ocean = green). Branch lengths are representative of nucleotide substitutions per site and the tree was midpoint rooted.

*N*_e_ trajectories showed significant phylogenetic signal in all time windows (p = 0.001; Table 2), indicating that closely related lineages tended to have more similar demographic histories than expected by chance. Kmult was lowest in the most recent window (50–100 kya, 0.259), peaked at 300–350 kya (0.668) and declined again in older windows. This is the inverse of the MANOVA result and suggests that older demographic history is more phylogenetically conserved, whereas recent *N*e shows the strongest regional divergence and the weakest phylogenetic structuring. This pattern is consistent with recent environmental drivers of regional divergence acting largely independently of shared ancestry.

**Table 2.**
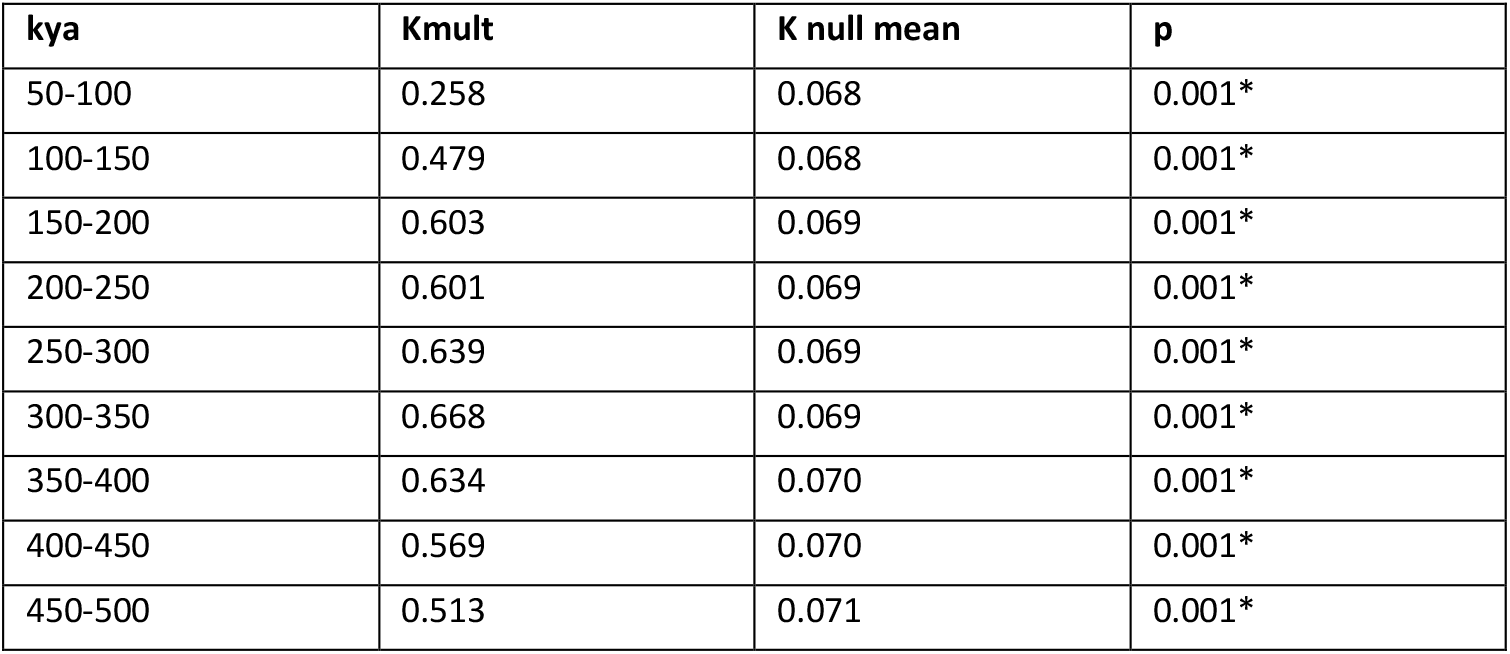
Kmult multivariate test of phylogenetic signal in effective population size. Values around 1 indicate a pattern consistent with Brownian motion, values below 1 indicate weaker similarity among related taxa than expected, and values above 1 indicate stronger similarity. Asterisk indicates statistical significance at *α* = 0.05.

| kya | Kmult | K null mean | p |
| --- | --- | --- | --- |
| 50-100 | 0.258 | 0.068 | 0.001* |
| 100-150 | 0.479 | 0.068 | 0.001* |
| 150-200 | 0.603 | 0.069 | 0.001* |
| 200-250 | 0.601 | 0.069 | 0.001* |
| 250-300 | 0.639 | 0.069 | 0.001* |
| 300-350 | 0.668 | 0.069 | 0.001* |
| 350-400 | 0.634 | 0.070 | 0.001* |
| 400-450 | 0.569 | 0.070 | 0.001* |
| 450-500 | 0.513 | 0.071 | 0.001* |

### 3.4 Phylogenetic multivariate analysis of variance (Phylogenetic MANOVA)

Phylogenetic MANOVA revealed significant differences in *N*_e_ trajectories among the three basins across sequential windows. Regional differences were strongest at 50–100 kya (F = 11.147, R² = 0.403) and 100–150 kya, and remained significant after maxT correction through 150–200 kya (p_maxT = 0.028; Table 3). The 200–250 kya window was marginally significant before correction (p_raw = 0.043) but not after (p_maxT = 0.181), with no significance in later windows. Southeast Asian lineages consistently showed the highest *N*_e_, Indian Ocean lineages the lowest, and Australian lineages intermediate values. Because Southeast Asia accounted for most between-group variance, we also contrasted it with the combined peripheral regions; this was significant through 50–200 kya after correction (p_maxT = 0.003, 0.001 and 0.033 for successive windows; Table 4) Because Southeast Asia contributed most of the between-group variance in the 3-group test, we also compared Southeast Asia versus all non-Southeast Asia lineages combined (AUS+IO). This contrast was significant through 50–200 kya after maxT correction (50–100 kya: p_maxT = 0.003; 100–150 kya: p_maxT = 0.001; 150–200 kya: p_maxT = 0.033; Table 4), supporting Southeast Asia as having consistently higher *N*_e_ compared to surrounding regions.

**Table 3.** Phylogenetic MANOVA across nine sequential time windows with permutation maxT correction for multiple testing. Three-group test (AUS/IO/SEA) with 999 permutations. Asterisk indicates statistical significance at *α* =

| kya | F | R2 | p (raw) | p (maxT) |
| --- | --- | --- | --- | --- |
| 50–100 | 11.147 | 0.403 | 0.001* | 0.002* |
| 100–150 | 13.053 | 0.442 | 0.001* | 0.002* |
| 150–200 | 5.818 | 0.261 | 0.002* | 0.028* |
| 200–250 | 3.214 | 0.163 | 0.043 | 0.181 |
| 250–300 | 2.377 | 0.126 | 0.095 | 0.325 |
| 300–350 | 2.056 | 0.111 | 0.142 | 0.408 |
| 350–400 | 1.738 | 0.095 | 0.175 | 0.494 |
| 400–450 | 1.473 | 0.082 | 0.217 | 0.577 |
| 450–500 | 1.277 | 0.072 | 0.260 | 0.644 |

**Table 4.** Phylogenetic MANOVA across nine sequential time windows with permutation maxT correction. Two-group test (AUS+IO versus SEA) with 999 permutations. Asterisk indicates statistical significance at =

| kya | F | R2 | p (raw) | p (maxT) |
| --- | --- | --- | --- | --- |
| 50–100 | 11.147 | 0.403 | 0.001* | 0.003* |
| 100–150 | 13.053 | 0.442 | 0.001* | 0.001* |
| 150–200 | 5.818 | 0.261 | 0.002* | 0.033* |
| 200–250 | 3.214 | 0.163 | 0.060 | 0.187 |
| 250–300 | 2.377 | 0.126 | 0.110 | 0.340 |
| 300–350 | 2.056 | 0.111 | 0.145 | 0.404 |
| 350–400 | 1.738 | 0.095 | 0.185 | 0.497 |
| 400–450 | 1.473 | 0.082 | 0.249 | 0.587 |
| 450–500 | 1.277 | 0.072 | 0.312 | 0.656 |

Among individual pairwise tests, Southeast Asia versus IO showed the strongest and most persistent signal, remaining significant after maxT correction through 150–200 kya (peak at 100–150 kya: F = 24.406, R2 = 0.670, p_maxT = 0.003). Southeast Asia versus Australia was significant in the earlier windows (50–100 kya: p_maxT = 0.004; 100–150 kya: p_maxT = 0.040) but not later. In contrast, Australia versus Indian Ocean showed a delayed pattern, with no significance at 50–100 kya (p_maxT = 0.739) but significant differences emerging at 100–150 kya and 150–200 kya (both p_maxT < 0.035).

### 3.5 Phylogenetic Least Squares (PGLS) regression of *N*_e_ and range size

The geographic range size of each lineage extracted from their Extent of Occurrence (EOO) ranged from 8.99e5km^2^ to 7.36e6km^2^ in Indian Ocean, 2.29e6km^2^ to 5.50e6km^2^ in Southeast Asia, and 7.70e4km^2^ to 5.83e6km^2^ in Australia. Southeast Asian lineages had the largest range size on average with a mean of 4.21E+06 km^2^ ± 8.82E+05, followed by IO with a mean EOO of 3.48E+06 km^2^ ± 3.54E+06. Australia had the lowest mean range size of 1.40E+06 km^2^ ± 1.10E+06.

Effective population size trajectories (50–500 kya) showed a significant positive relationship with range size when region was not included as a covariate (F = 7.576, R2 = 0.187, r = 0.432, p = 0.006; n = 35) (Supplementary Table S5). This is *N*_e_ ∼ log(EOO). This suggests that lineages with larger geographic ranges tend to have higher recent *N*_e_. However, when ocean basin was included as a covariate (*N_e_* ∼ region + log(EOO)), EOO was no longer significant (F = 1.086, R2 = 0.020, p = 0.321), while region remained strongly significant (F = 11.047, R2 = 0.408, p = 0.001). This result indicates that the apparent EOO–*N*_e_ relationship was driven by regional structure: Southeast Asian taxa have both larger ranges and higher *N*_e_ than Australian or Indian Ocean taxa, generating a correlation between EOO and *N*_e_ that disappears once regional identity is accounted for.

### 3.6 Population Diversity Statistics

Diversity statistics were estimated for the genetically distinct populations identified by PCA, fastMIXTURE and tree topology (Supplementary Table S6), and compared among basins for the 49 populations with five or more individuals (Figure 7). Nucleotide diversity (*π*) and observed heterozygosity (H_o_) values followed similar patterns for each ocean basin, with highest means in Australia, followed by Southeast Asia and Indian Ocean with the lowest. The population with both the highest nucleotide diversity (*π* = 0.245) and observed heterozygosity (H_o_ = 0.222) was the *H. kingii* population in West Kimberley, Australia.

**Figure 7.**
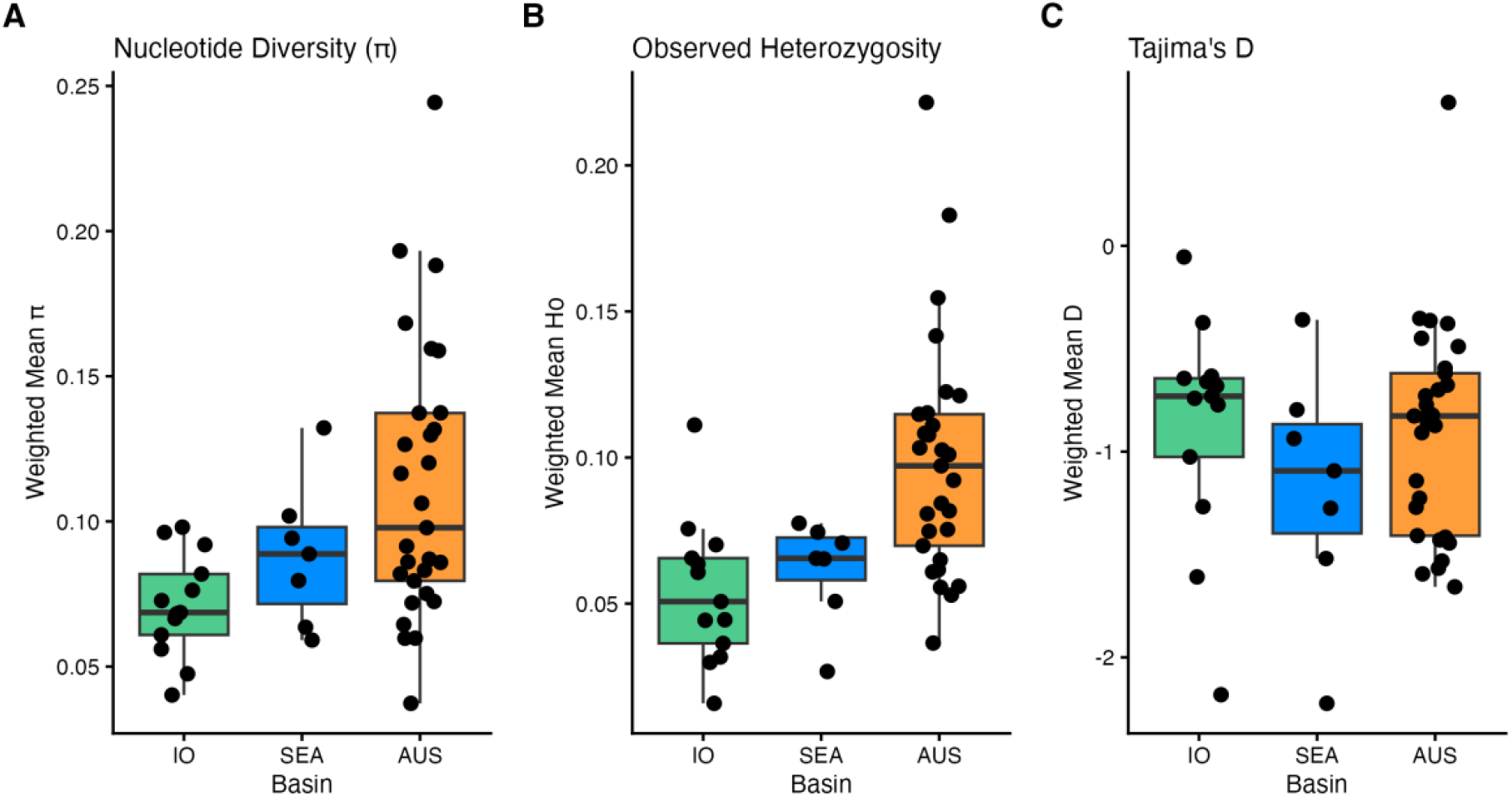
Population statistics for (A) nucleotide diversity, (B) observed Heterozygosity and (C) Tajima’s D (C) of genetically distinct populations (n >= 5) calculated in snpR. The weighted means of each statistic accounted for differences in sample size.

All populations, except *H. major* in New Caledonia, had a negative Tajima’s D statistic resulting from an excess of rare alleles indicating recent population expansion across all basins. Only *H. major* in New Caledonia had a positive D statistic (0.631) indicating recent population bottleneck. Southeast Asia had the lowest mean D statistic suggesting the strongest signal of recent expansion among basins. The weakest signal of recent population expansion was observed in the Indian Ocean mean D statistic. However, the *H. ornatus* population from Andaman Sea did not follow this general trend with one of lowest D statistics (-2.158).

## 4 Discussion

Quantifying variation in the demographic trajectories of co-distributed taxa is central to anticipating how populations will vary in their response to ongoing climate change, and where such comparisons span biogeographic regions they can also illuminate the origins of biodiversity hotspots. Genome-wide SNP data for more than 650 individual sea snakes recovered clear splits among closely related lineages separated across the Indian Ocean (IO), Southeast Asian (SEA) and Australian (AUS) biogeographic regions, consistent with previous inferences of Pleistocene isolation and vicariant diversification in sea snakes and other marine taxa (Ludt and Rocha, 2015; Cowman and Belwood, 2013; García-Jiménez et al., 2025; Hsu et al., 2025; Tang et al., 2025, Ukuwela et al., 2016). The concordance justified using coalescent analyses of whole genomes to test whether *N*_e_ trajectories are structured more strongly by regional history than by phylogeny or range size. Southeast Asian lineages consistently showed the largest historical *N*_e_ and the shallowest bottlenecks compared to those from Australia and the Indian Ocean. These results are discussed below in the context of biogeographic processes shaping the IAA hotspot and their implications for conservation prioritisation under ongoing climate change.

### 4.1 Demographic support for the Indo-Australian Archipelago as a centre of sea snake diversity

*N*_e_ trajectories in the three basins diverge between approximately 50 and 200 kya, consistent with the geographic structuring revealed by the SNP data and with Pleistocene isolation among basins. Southeast Asian lineages consistently show the largest historical *N*_e_ and the shallowest bottlenecks, supporting a role for the region as a long-term centre of population persistence; the trend is most pronounced over the last 50–200 kya and attenuates in older periods. The inverse pattern is observed in the multivariate analyses of phylogenetic signal, in which the influence of shared ancestry in shaping *N*_e_ is weakest in the most recent time intervals and increases toward older time periods. Together, these results suggest that the demographic patterns of co-distributed sea snakes have been shaped more strongly by regional history than by conserved phylogenetic responses. Lineage-specific traits relating to dispersal propensity and eco-morphological specialisation must also contribute, and quantifying their effects should be a priority for future studies. However, many of the same ecomorphs have evolved convergently in each of the three regions (Sherratt et al., 2018), suggesting that regional differences in *N*_e_ are not strongly influenced by among-basin biases in lineage traits.

The elevated and relatively stable *N*_e_ of sea snakes in Southeast Asia is consistent with the well documented role of this region as a refugium for shallow marine taxa during Pleistocene glacial cycles. Under the centre of refugium hypothesis, relic populations persisted in stable habitats through repeated the Pleistocene sea-level fluctuations, and our results might suggest that proximity to persistent shallow-water habitat was important in preserving genetic diversity. Substructure between the northeast and southwest of the Sunda Shelf is further evidence of the refugial history where a land bridge during glacial lows restricted gene flow and facilitating vicariant diversification (Ludt and Rocha, 2015). Here deep basins including the South China Sea, Sulu Sea, Celebes Sea to the north-east of Sunda and the Andaman Sea to the southwest supported shallow water habitat during the lowest sea levels of the Pleistocene (Bellwood and Hughes, 2001; Woodruff, 2003; Ludt and Rocha, 2015).

Fluctuating *N*_e_ in the peripheral Indian Ocean and Australian regions suggests a history of population bottlenecks and expansions as low sea-levels exposed seabeds (Pellissier et al., 2014), while negative Tajima’s *D*) in all three ocean basins indicates contemporary expansion in all regions. Australia also had a longer history of diversification, evidenced_by deeper phylogenetic splits, and the nesting of Southeast Asian clades within Australian species suggests dispersal from Australia into Southeast Asia. Diversity from the peripheral region has therefore contributed to the accumulation of high diversity within the IAA hotspot, consistent with models of hotspot formation in which peripheral regions act as refuges for older taxa (Briggs, 1999; Bellwood et al., 2012). Our results are compatible both with the ‘centre of’ hypotheses and with more recent concepts of hotspots that shift through evolutionary time, under which the IAA hotspot formed around 1–2 Ma, coinciding with the sea snake radiation (Huang et al., 2025).

### 4.2 Implications for sea snake conservation

The deepest phylogenetic splits were recovered between the three ocean regions (IO, SEA, AUS), which therefore represent distinct evolutionary and conservation management units. Finer-scale patterns of geographic genetic structure were found to vary among basins (diversity and expansion metrics) with important implications for spatial planning of marine protected areas, spatial scale of threats, and recovery from local declines. Not all of these geographic patterns reflect intermittent historical barriers (glacial cycles), many are consistent with contemporary oceanographic features that may also contribute to spatial pattern of genetic diversity within the regions. Region-specific findings and their implications are discussed below.

Clear regional structuring of demographic histories indicates that each biogeographic region represents a distinct component of sea snake diversity shaped by distinct environmental histories. This suggests the need for region-specific conservation strategies, in addition to approaches based on species-level conservation status. A further concern is the concentration of data-deficient species in Southeast Asia and the Indian Ocean. The Indian Ocean region is of particular concern: its lineages are demographically distinct and may retain unique genetic diversity despite consistently lower *N*_e_, but the region also shows greater demographic vulnerability and limited historical refugial capacity, and is therefore likely to be more sensitive to the intense habitat loss and fishing pressure it currently faces. Targeted monitoring and protection are warranted. Our findings also have important implications for marine conservation in the Indo-Australian Archipelago under ongoing environmental change. Signals of long-term demographic stability in Southeast Asian lineages of sea snakes highlight the importance of structurally complex, historically stable shallow marine habitats for sustaining populations. This historical context contrasts sharply with contemporary habitat loss and intense fishing pressure.

Contemporary genetic diversity was highest in Australia, despite the larger ranges of Southeast Asian lineages. This may partly reflect denser Australian sampling, since pooling individuals across localities can inflate nucleotide diversity. However, the pattern persisted when sample sizes were equalised, and more likely reflects recent expansion as the continental shelf was reflooded, consistent with rising Australian Ne towards 50 kya.

Genetic structure within regions indicates that finer-scale management aligned to contemporary biogeographic features is also required. For Australian species, this was the first population genetic assessment of *H. peronii* and *H. ocellatus* and the first of any SNP-based population genetic analysis for Southeast Asia and Indian Ocean sea snakes, covering seven species here (*H. curtus*, *H. ornatus*, *H. schistosus*, *H. spiralis*, *H. cyanocinctus*, *H. fasciatus*, *H. atriceps*). Within the Australia-West Pacific biogeographic region, substructure differed among species in degree, but was most prominent between the west (West Australia and Timor Sea) and the east (North Australia, East Australia and New Caledonia), either side of the Kimberley. This region acts as a biogeographic barrier for other shallow-water marine taxa due to its complex geology and large tidal movements (Wilson, 2014). In the Indian Ocean, common patterns of broad scale biogeographic structuring among species were difficult to assess due to lack of sampling depth and localities. Sufficient sampling is required to test whether the weak structuring follows an isolation-by-distance pattern and to assess the connectivity of these populations. In Southeast Asia, we observed distinct structure across the region’s complex geology including to the northeast and southwest of the Sunda Shelf, which was exposed during Pleistocene glacial periods. Further dense sampling in this hotspot region is required to explore the influence of specific/contemporary oceanographic features on sea snake diversity.

### 4.3 Unrecognised taxonomic diversity and sampling priorities

The depth of intraspecific structure recovered between basins, particularly in *H. curtus* and *H. ornatus*, suggests the presence of unrecognised species. Phylogeographic structure among basin-endemic lineages within these species is comparable in scale to that between nominal sister species such as *H. fasciatus* and *H. atriceps*. These lineages are not cryptic: several appear morphologically diagnosable, and *H. curtus* was treated as two species for much of its taxonomic history. Combined with evidence that these populations have been shaped by distinct demographic histories, this indicates that taxonomic revision is needed to describe unrecognised diversity within widely distributed species.

Resolving this diversity will require further sampling. The present work substantially advanced knowledge of demographic history and population trends for a third of all sea snake species, but it primarily examined patterns between distant localities along the Sunda, Sahul and Indian Ocean shelves, and lacks resolution at finer spatial scales in several key regions. Sampling gaps in the Indian Ocean are of particular concern for India, where population declines and destructive fishing practices are documented (Elfes et al., 2013; Udyawer et al., 2018). Southeast Asia is under-sampled across its complex oceanography, including the Philippines and Japan, where sea snake diversity remains poorly described. In the Australia-West Pacific, Papua New Guinea and the western Pacific islands are a conspicuous sampling gap. Targeting these regions should be a priority for future work.

## Data availability Statement

All data generated for this project has been made publicly available on NCBI BioProject x (WGS) and y (Dartseq). Scripts and data are available on GitHub (https://github.com/AmeliaPointon/global-sea-snakes-phylogenomics).

## Animal Ethics and Collection Permits

All sampling was in accordance with methods approved by the University of Adelaide’s Animal Ethics committee (permit 34903), James Cook University (permit A2310), Charles Darwin University Animal Ethics (permit A18029). Collection permits were granted from the Western Australian Department of Biodiversity, Conservation and Attractions (FO25000393-2, DPAW08-000725-1, DoEE E2017-01290, and Fisheries collection exemption 2527 from the Department of Primary Industries and Regional Development), Department of Wildlife and Conservation of Sri Lanka (WL/3/2/1/14/12), and the New Caledonia Province Sud (3252-2017/ARR/DENV).

## Acknowledgements

This work was supported with supercomputing resources provided by the Phoenix HPC service at Adelaide University. The project received funding by the Australian Research Council Discovery Grant (DP180101688), the Australian Government’s Our Marine Parks Grant Round 3 (4-H0MJUFC), and the Australian Government’s National Environmental Science Program. We also thank Claire Gorian, Kanishka Ukuwela, Abdur Razzaq Sarkar, Arne Rasmussen and Isaac Rossetto for sample collection. We thank Matthew Ford and Shannon Coppersmith for assistance with lab work, which was conducted at the South Australian Museum’s SARFMEE facility. We thank the following museums for access to tissue samples: Field Museum of Natural History (FMNH), Museum and Art Gallery of the Northern Territory (MAGNT), Museum Zoologicum Bororiense (MZB), Queensland Museum (QM), South Australian Museum (SAMA) and Western Australian Museum (WAM).

## Author contributions

A.R.P, A.J.L. and K.L.S. designed the research idea. K.L.S., V.U., J.H.N., V.O.G., A.R.P., B.B., B.F., and C.G. collected samples. A.J.L ran the MSMC2 analysis. V.U. and V.O.G produced Extent Of Occurrence datasets. A.R.P. and J.H.N. conducted the laboratory work. A.R.P. conducted all other analyses and led the writing. All authors contributed to interpreting the results and reviewing and editing the manuscript.

## Supplementary

Supplementary data tables are available in supporting documents.

**Supplementary Figure 1.**
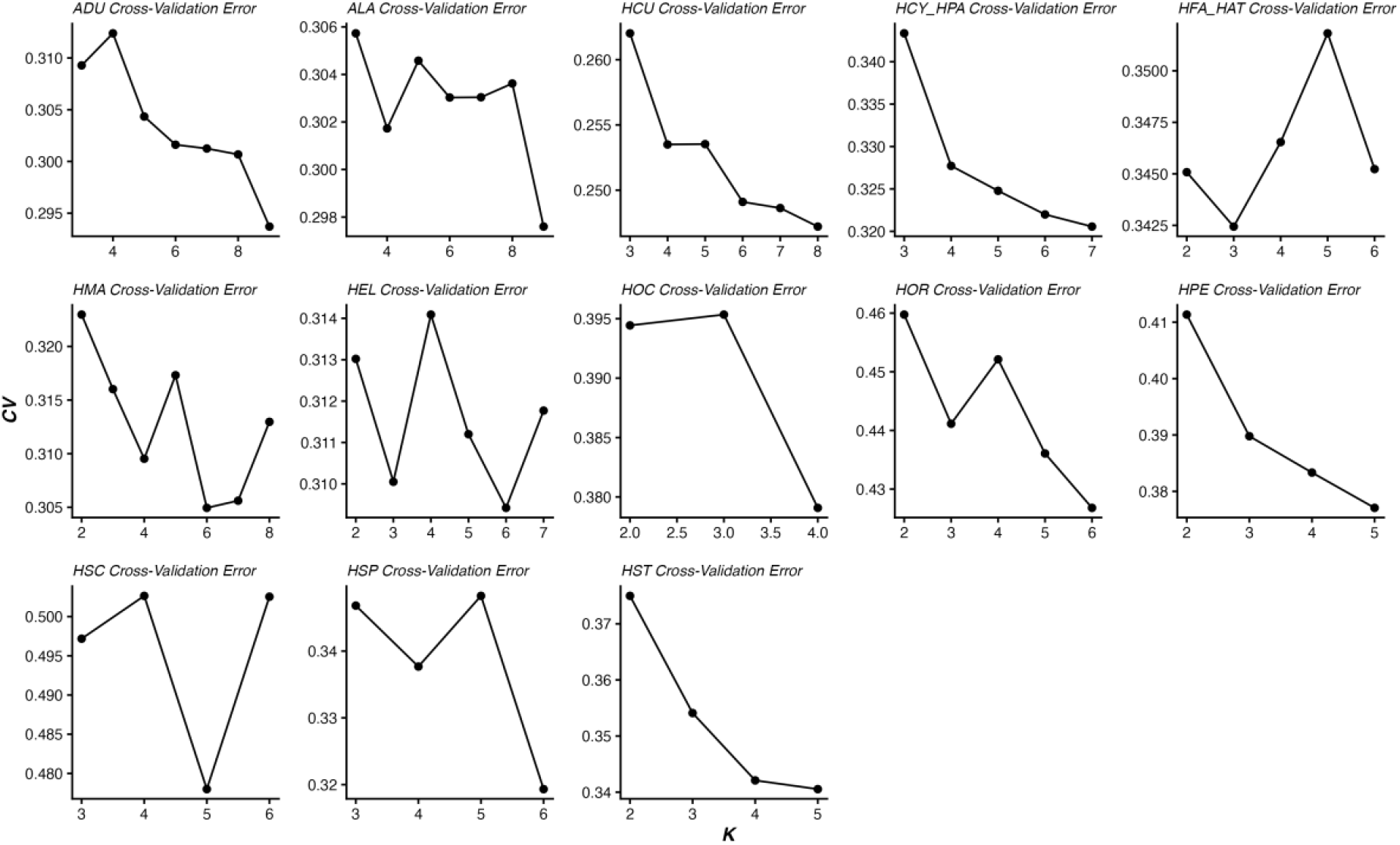
Admixture Cross-validation Error across *K* number of ancestry clusters for each species analysed in fastMIXTURE.

**Supplementary Figure 2.**
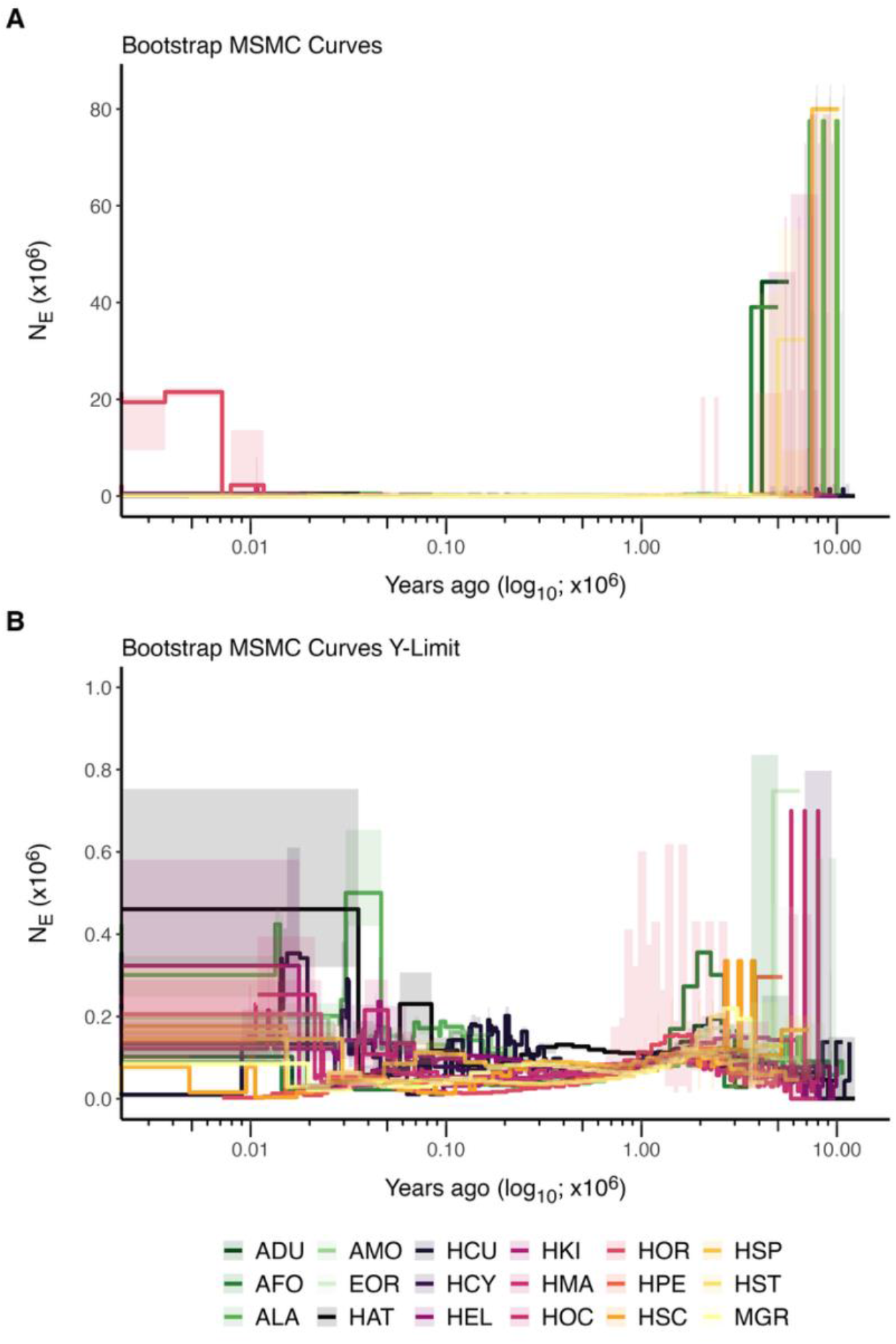
Raw effective population size (*N*_e_) curves for each individual sea snake coloured by species. Minimum and maximum bootstrap values in shaded areas. (A) Raw *N*_e_ data are shown with no axis limitations from 10MYA to present day. (B) Raw *N*_e_ data are shown with a maximum limit of 1e6 applied to the y-axis.

